# Stimuli−Expressed Protein Qubits for Quantum Quantification of Heavy Metals in Natural Water

**DOI:** 10.64898/2025.12.04.692285

**Authors:** Wenhao Shan, Fujin Lv, Wei Liu, Bin Li, Yingying Wang, Xianli Gong, Fulin Zhu, Bingchen Zeng, Tiantian Man, Qiuyue Wu, Shengyuan Deng

**Affiliations:** School of Environmental and Biological Engineering, Nanjing University of Science and Technology, Nanjing 210094, China; School of Mechanical Engineering, Nanjing University of Science and Technology, Nanjing 210094, China

**Keywords:** Environmental heavy metal pollution, Quantum biosensing system, CadC−T7 gene circuit, Magnetically modulated fluorescent protein, Lock-in analysis

## Abstract

Environmental heavy metal pollution poses a severe threat to ecological security and human health, necessitating the development of highly sensitive, specific, and rapid detection technologies. Traditional fluorescent biosensors are often constrained by high background interference and limited sensitivity, while single-target detection systems struggle to meet the demands of complex environmental monitoring. This study constructs a novel magnetically modulated photonic quantum biosensing system based on the CadC−T7 gene circuit, replacing the traditional fluorescent reporter gene EGFP with the magnetically modulated fluorescent protein MagLOV - a quantum biomaterial. To enhance its magnetic modulation performance, the mutant with the highest lock-in amplitude during the directed evolution (R10, MagLOV 2) was selected; Simultaneously, systematic parameter optimization was conducted to determine the optimal magnetic modulation frequency (0.05 Hz), the heavy metal stimulation duration (the 2-hour group exhibited the most significant intra-group variation), and the bacterial concentration at the onset of the heavy metal stimulation (controlled at OD_600_ = 1 to prevent fluorescence quenching due to culture turbidity). The system demonstrates excellent multi-analyte detection capabilities for lead chloride, lead nitrate, and cadmium chloride across a concentration range of 10^−8^ to 10^−4^ M. Compared to conventional steady-state photometric methods based on EGFP and MagLOV 2, the lock-in analysis of MagLOV 2 achieves detection sensitivities enhanced by 1∼2 orders of magnitude, with detection limits as low as 2.4 nM for cadmium chloride, 4.8 nM for lead nitrate, and 9.9 nM for lead chloride. Langmuir isotherm fitting results demonstrate high quantitative accuracy (*R*^2^ > 0.950), with particularly outstanding fitting performance for cadmium chloride (*R*^2^ = 0.999) and lead nitrate (*R*^2^ = 0.995).

## 1. INTRODUCTION

Heavy metals, such as cadmium and lead, characterized by high toxicity, persistent nature, and bioaccumulative effects, have become critical pollutants requiring urgent management in soil and aquatic environments. These substances can accumulate progressively through the food chain, causing neurological damage, organ dysfunction, and even cancer in humans, posing a severe threat to ecosystem stability and human health.^[1−5]^ Therefore, it is crucial to develop direct and highly sensitive tools for determining heavy metal concentrations in the environment. Currently, various methods such as atomic absorption spectroscopy (AAS), ultraviolet-visible spectroscopy (UV), X-ray fluorescence (XRF), and inductively coupled plasma mass spectrometry (ICP−MS) have been employed for the determination of heavy metal ions.^[6−9]^ These methods exhibit high sensitivity and selectivity, but they require expensive instruments, complex operating procedures, and lengthy detection times. Consequently, they are neither cost-effective nor suitable for routine heavy metal analysis.^[6,10−13]^ Accordingly, developing low-cost heavy metal detection technologies that combine high specificity, high sensitivity, and practical convenience has become an urgent task in the field of environmental monitoring. Biosensors, which detect target substances through the specific interaction between their biological recognition elements and target analytes, possess the core advantages of high specificity, high sensitivity, low cost, and practical convenience, thus having emerged as a highly promising and important development direction in this field.^[14−18]^ Among these, microbial gene signaling pathway sensors based on fluorescence reporting systems are widely applied. They achieve visual and quantitative detection of target substances by driving reporter protein expression through target-responsive elements.^[19−23]^ Such systems predominantly employ traditional fluorescent proteins like green fluorescent protein (GFP, EGFP) as reporter units. However, the fluorescence signals from these proteins are susceptible to interference from biological autofluorescence and light scattering, resulting in a low signal-to-background ratio (SBR). This limitation constrains improvements in detection sensitivity and other performance metrics.^[24−27]^

To overcome the bottleneck of high background fluorescence noise and significant signal drift, quantum nanomaterials (such as fluorescent nanodiamonds, FND) have emerged.^[28−31]^ FND leverages the quantum magnetic response effect of nitrogen-vacancy (NV⁻) centers to achieve magnetic field-fluorescence synchronous modulation under an applied magnetic field. This transforms the emission mode from traditional steady-state emission prone to photobleaching or random scintillation into a frequency-stable, waveform-regular light-switching mode. Consequently, lock-in detection technology can filter out non-periodic biological spontaneous fluorescence and scattering noise, The theoretical sensitivity is enhanced by 2∼3 orders of magnitude compared to steady-state imaging.^[32]^ Lock-in detection demonstrates significant value in biological detection, enabling precise differentiation of fluorescently labeled molecules or cells within tissues or environments with high background signals - where background signals often originate from the spontaneous fluorescence of tissues and neighboring molecules.^[33,34]^ To address this core challenge, relevant research has quantified the key evaluation metric - SBR, providing a clear basis for assessing the application of the technology.^[35]^ However, when applied to living models, FND’s inherent limitations become immediately apparent: it requires chemical conjugation with antibodies/peptides for localization, exhibits significant batch-to-batch variability; residual carboxyl-PEG or cationic polymers on surfaces can trigger immune responses; and its manufacturing costs are high, necessitating high-temperature/high-pressure synthesis plus electron irradiation. The emergence of magnetically modulated fluorescent proteins (MagLOV) offers a novel technical solution to these challenges. Based on the radical pair mechanism (RPM), MagLOV exhibits periodic bright-dark fluctuations in fluorescence intensity under external magnetic field control. It combines quantum-level magnetic response sensitivity with prokaryotic *in situ* expression capability, eliminating the need for complex chemical modifications.^[36]^ Compared to FND’s reliance on chemical conjugation, MagLOV’s *in situ* expression capability preserves high sensitivity while avoiding drawbacks such as insufficient biocompatibility and complex operation. Its low cost and scalability, coupled with the ability to target subcellular compartments like synapses and mitochondria, achieves a unified approach of high sensitivity, high adaptability, and low cost for heavy metal detection.

Against this backdrop, this study successfully integrated three core advantages: quantum-level magnetic response sensitivity, prokaryotic *in situ* expression characteristics, and outstanding biocompatibility, to construct a novel quantum bio-detection system featuring MagLOV 2 as its signal reporting unit. This breakthrough effectively overcomes the dual technical limitations that have constrained the application of traditional fluorescent proteins with low SBRs and the limitations of quantum nanomaterials like fluorescent nanodiamonds (FNDs) in *in vivo* applications (Figure 1, prototype shown in Figure S1). The study has successfully established an *in situ* expression system for MagLOV 2 based on the CadC−T7 gene circuit,^[37,38]^ confirming that MagLOV 2 can be stably expressed with uniform distribution in *E. coli*. Its spectral characteristics remain consistent with purified proteins, while exhibiting no significant toxicity to host cells and demonstrating excellent biocompatibility. Through systematic optimization of key detection parameters, including magnetic modulation frequency (optimal value: 0.05 Hz) and induction time (optimal value: 2 h), the system’s SBR and detection sensitivity for cadmium and lead ions were significantly enhanced. In gradient detection of 10^−1^^1^∼10^−4^ M concentrations of lead chloride, lead nitrate, and cadmium chloride, MagLOV 2 demonstrated significant performance advantages in magnetically modulated photonic qubit lock-in analysis. Its detection limit was reduced by 1∼2 orders of magnitude (down to 2.4∼9.9 nM) compared to EGFP and MagLOV 2 steady-state fluorescence photometry. Langmuir isotherm fitting *R*²values exceeded 0.95, markedly superior to the 0.87∼0.97 ranges of the latter two methods. Particularly within the 10^−8^∼10^−4^ M concentration range, SBR enhancement was more pronounced, with optical quantum bit signal discrimination far surpassing both traditional fluorescence detection approaches. This fully demonstrates the signal amplification and anti-interference advantages of magnetically modulated optical quantum bit lock-in technology. This method achieves detection limits at the trace level, providing novel technical support and a reliable methodological reference for rapid trace detection of heavy metals in environmental samples.

**Figure 1.**
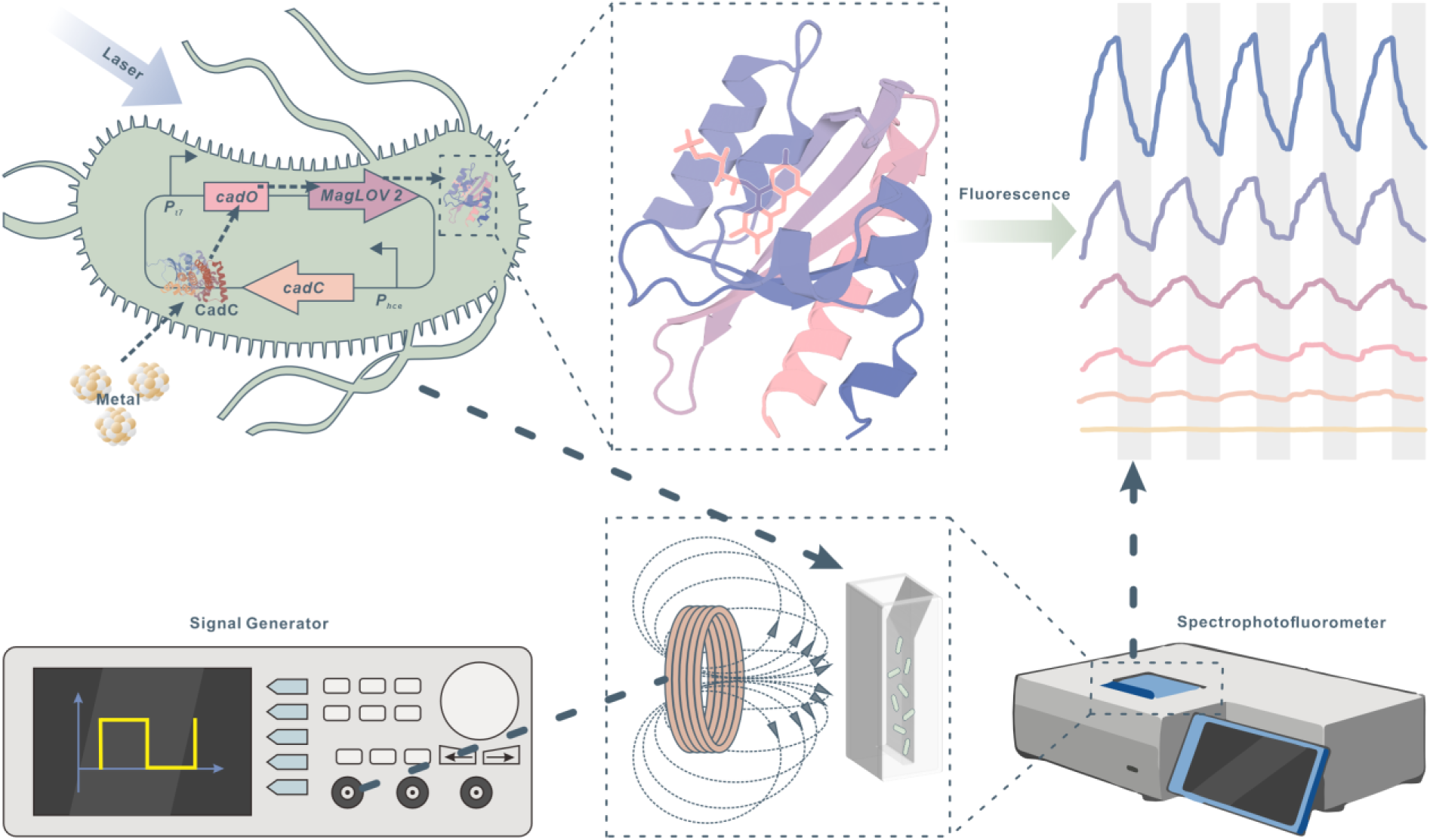
Schematic of *E. coli* expressing magnetically modulated fluorescent protein for detecting heavy metal ions. Heavy metal-responsive genes within the CadC−T7 circuit continuously express the response protein CadC during bacterial growth. Upon reaching a certain expression threshold, CadC is activated by specific heavy metal ions and acts on the corresponding cadO operon, thereby unlocking expression of the downstream reporter gene *MagLOV*. An applied periodic magnetic field (gray) modulates the 490 nm green fluorescence emitted by MagLOV upon excitation by 460 nm blue light, inducing coherent on-off transitions. The coherent fluorescence signal detected by a spectrophotofluorometer can be further processed using lock-in amplitude signal processing.

## 2. Magneto-Modulated Fluorescence of MagLOV in vitro

To investigate the optical properties and magnetic response mechanism of the magnetically modulated fluorescent protein MagLOV, we conducted systematic *in vitro* experiments with purified MagLOV and flavin mononucleotide (FMN) in PBS buffer. Vector construction is detailed in the session S1.3 of the Supporting Information (SI), protein expression and purification in S1.4, and magnetic regulation and detection in S1.5. As shown in Figure 2, the fluorescence spectrum overlay of MagLOV and FMN reveals a characteristic strong fluorescence peak at approximately 490 nm for MagLOV, while FMN exhibits its peak at around 525 nm. This spectral overlay analysis confirms that the potential spurious peak in the MagLOV spectrum originates from FMN (Figure 2a). This eliminates potential interference from cofactor spectra, ensuring that the observed fluorescence changes stem from the protein’s intrinsic magnetic regulation effect and enabling precise analysis of MagLOV’s magnetic response characteristics. Under a 10-mT periodic magnetic field, the fluorescence intensity of 2 μM FMN showed no significant periodic fluctuations. This result directly demonstrates that free FMN lacks magnetic modulation capability, thereby ruling out the possibility of cofactor FMN influencing MagLOV’s magnetic response characteristics. It further indicates that MagLOV’s magnetic response properties originate from its protein structure itself (Figure 2b). In contrast, the fluorescence intensity of 2 μM MagLOV exhibited regular fluctuations in response to the periodic magnetic field. However, the amplitude of the magnetic response fluctuations was relatively mild when comparing signal magnitudes (Figure 2c). Compared to the 2 μM concentration group, the fluorescence intensity fluctuations of 10 μM MagLOV were more pronounced, with significantly increased magnetic modulation signal amplitude and more intense periodic bright-dark state transitions (Figure 2d). This result indicates that MagLOV’s magnetic modulation effect exhibits concentration dependence, with higher concentrations demonstrating superior fluorescence response characteristics under magnetic regulation. This concentration dependence manifests as a positive correlation between MagLOV fluorescence modulation amplitude and protein concentration, providing a critical application basis for quantitative analysis scenarios such as heavy metal ion detection. By leveraging the stimulation mechanism of heavy metals on MagLOV expression, changes in the lock-in amplitude of its magnetically modulated photonic qubits can reflect MagLOV expression levels, thereby enabling quantum-level quantitative detection of heavy metal concentrations.

**Figure 2.**
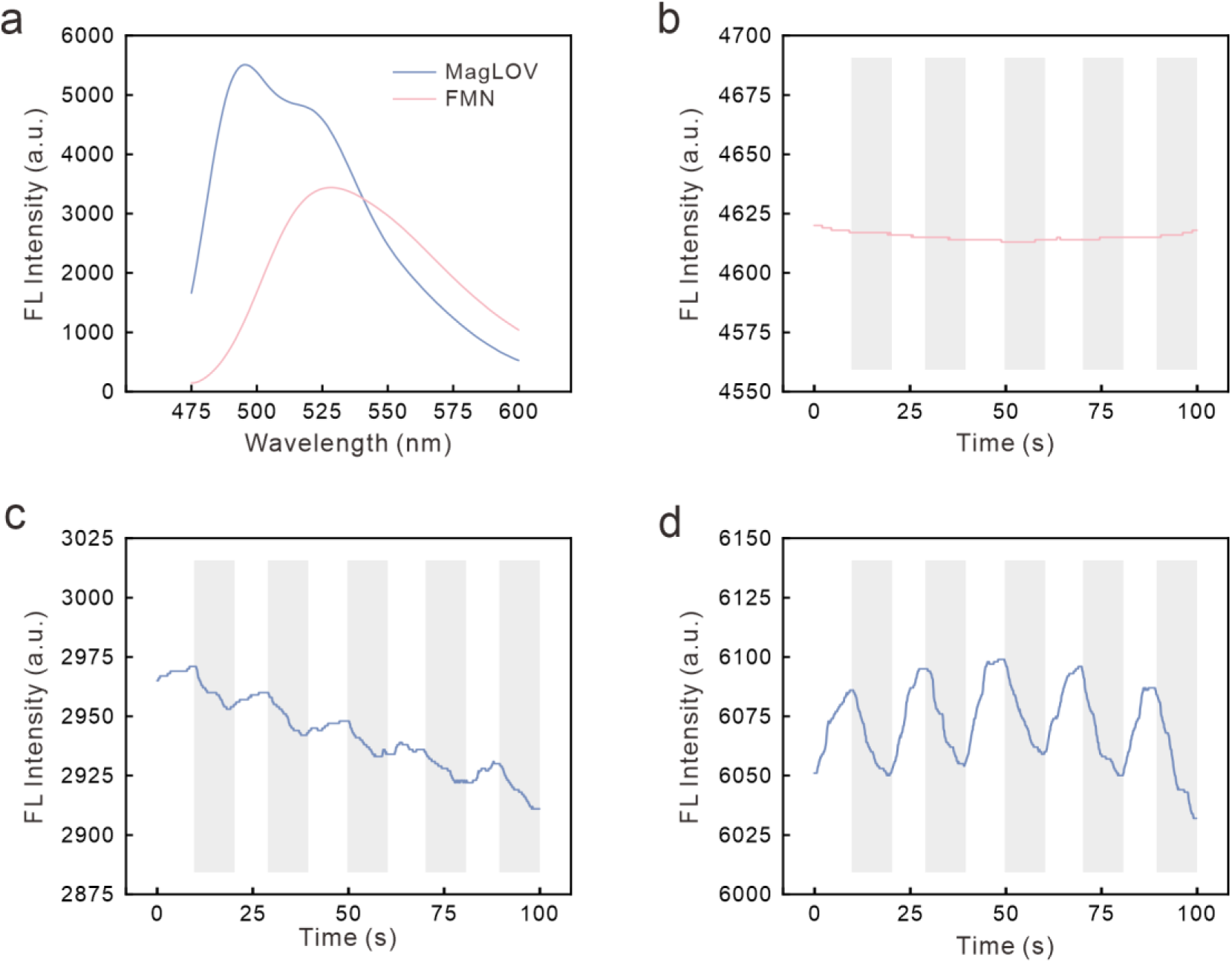
In vitro spectra of purified MagLOV and FMN in PBS buffer, along with magnetic modulation time-domain fluorescence spectra at a magnetic field strength of 10 mT. **a)** Overlay of fluorescence spectra for MagLOV and FMN; **b)** Magnetically modulated fluorescence time-domain spectrum of 2 μM FMN, **c)** 2 μM MagLOV, and **d)** 10 μM MagLOV.

## 3. Magneto-Modulated Fluorescence of MagLOV in vivo

The *in vivo* fluorescence characteristics of MagLOV expressed in the strain are consistent with its purified *in vitro* fluorescence properties. MagLOV-induced BL21(DE3) exhibits a distinct characteristic fluorescence peak at approximately 490 nm, with significantly higher fluorescence intensity than uninduced and wild-type strains (Figure 3d). Confocal microscopy visualization provides visual confirmation fully consistent with the spectral data. The field of view for induced MagLOV-expressing BL21(DE3) shows abundant green fluorescence signals (Figure 3a). In contrast, the field of view for uninduced MagLOV-expressing BL21(DE3) exhibits only faint, minimal fluorescence (Figure 3b). The field of view for wild-type BL21(DE3) showed almost no fluorescence signal (Figure 3c). This synergistic evidence from quantitative spectral analysis and qualitative microscopic imaging fully demonstrates that MagLOV achieves highly efficient, inducible-specific expression in BL21(DE3), with its fluorescent properties fully preserved in the complex *in vivo* environment. This lays a crucial experimental foundation for subsequent development of an *in vivo* heavy metal ion biosensing detection system using this engineered *E. coli* strain, addressing both the *in situ* expression and functional efficacy of the protein within the bacterial host.

**Figure 3.**
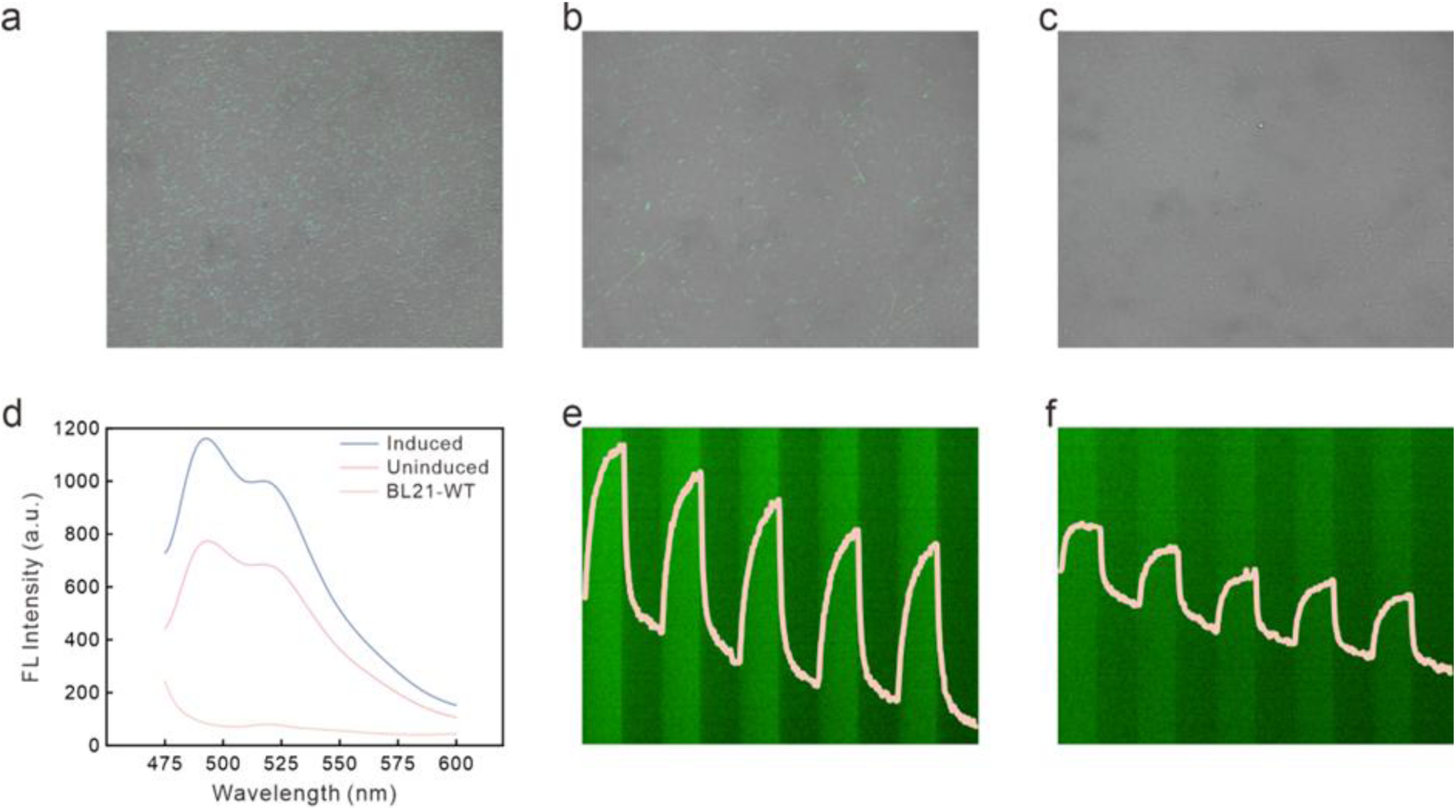
Confocal microscopy visualization field of view for **a)** induced MagLOV BL21(DE3); **b)** uninduced MagLOV BL21(DE3); **c)** wild-type BL21(DE3); and **d)** Spectral diagrams of induced MagLOV-expressing, uninduced MagLOV-expressing and wild-type BL21(DE3). Magnetically modulated fluorescence blinking profile of MagLOV-expressing BL21(DE3) at **e)** OD_600_ = 2.0 and **f)** OD_600_ = 0.5 obtained by Z-slice processing via Show Slice View function of NIS-Elements software.

First observation of the magnetically modulated fluorescence blinking of MagLOV was achieved by magnification via Ti2-U inverted FL microscope (Nikon, Japan) and capture using Nikon A1 camera (LFOV, 1024×1024 pixel2, Nikon, Japan). Utilizing the Show Slice View function in NIS-Elements software, we performed Z-slice processing on imaging videos of MagLOV-expressing BL21(DE3) at different concentrations, enabling the precise acquisition of Z-axis profiles of this magnetically modulated fluorescence blinking. For MagLOV-expressing BL21(DE3) at OD_600_ = 2.0 (Figure 3e), the fluorescence blinking profile exhibited more pronounced amplitude fluctuations - a feature directly attributed to the enhanced magnetically modulated signal response driven by the aggregation of MagLOV at this higher *E. coli* concentration. In contrast, the blinking profile of cultures at OD_600_ = 0.5 displayed relatively subdued fluctuations (Figure 3f), consistent with the lower molecular density of MagLOV at this reduced *E. coli* concentration.

To establish a quantitative relationship between the magnetically modulated fluorescence performance of MagLOV in *E. coli* and *E. coli* concentration, thereby providing key performance parameters for subsequent development of magnetic modulation biosensing systems for heavy metal ions, we conducted a systematic analysis of magnetic modulation characteristics in MagLOV-expressing BL21(DE3) at varying concentrations (characterized by OD_600_) under a magnetic field strength of 10 mT. The time-domain profiles of magnetic modulation for this engineered strain at different concentrations clearly demonstrate that as the OD_600_ of BL21(DE3) gradually increased from 0.01 to 2.00, the fluctuation amplitude of the magnetically modulated photonic qubit exhibited a distinctly stepwise enhancement trend. Signal waveforms across concentration groups showed clear distinguishability in periodicity and amplitude, directly reflecting the concentration-dependent regulation of the magnetically modulated photonic qubit signal (Figure 4a). However, it should be noted that further increases in *E. coli* concentration led to ineffective fluorescence detection due to optical scattering and absorption properties of the bacterial suspension, resulting in a continuous decline in signal intensity. The lock-in amplitude output diagram of the magnetically modulated optical quantum bit quantitatively demonstrates a significant positive correlation between lock-in amplitude and bacterial OD_600_, clearly quantifying the linear response relationship between bacterial concentration and optical quantum bit lock-in amplitude (Figure 4b). Based on the above analysis, subsequent experiments must strictly maintain *E. coli*’s detection concentrations at OD_600_ = 1 to prevent detection failure caused by excessive bacterial density while ensuring stable and controllable magnetic-modulated photonic qubit lock-in signals.

**Figure 4.**
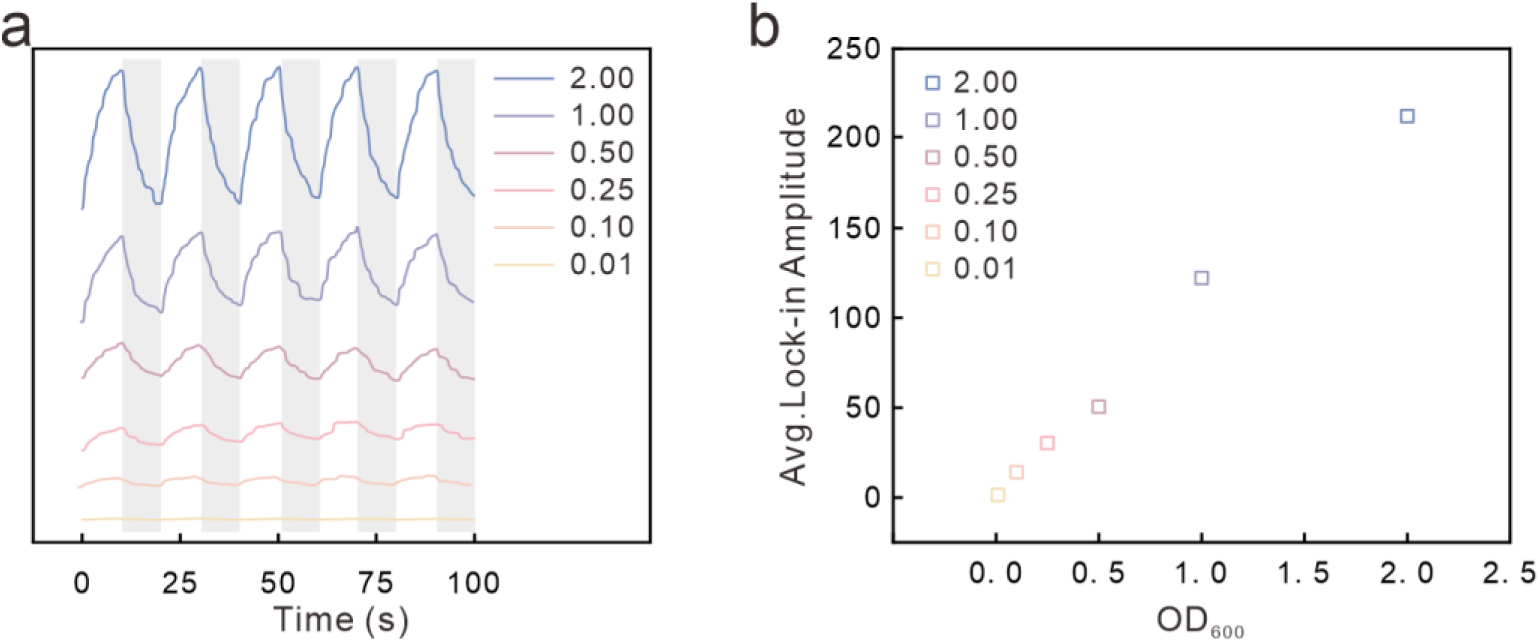
**a)** Time-domain profile of magnetically modulated photonic qubits from MagLOV-expressing BL21(DE3) at a magnetic field strength of 10 mT under several concentration conditions, and **b)** lock-in amplitude output of magnetically modulated photonic qubits obtained via fast Fourier transform.

## 4. Selection of Directed Evolution Mutants

To select MagLOV variants with superior magnetic modulation performance that meet the high-sensitivity reporter gene requirements for subsequent heavy metal ion detection, we conducted reproducibility analyses of magnetic modulation characteristics in representative MagLOV-induced BL21(DE3) strains from previously reported directed evolution processes.^[36]^ Under a magnetic field strength of 10 mT, the time-domain profiles of magnetic modulation for MagLOV mutants across different rounds expressed in BL21(DE3) revealed significant variations in signal amplitude: the R10 strain exhibited markedly higher modulation amplitude than other rounds (Figure 5a). The lock-in amplitude plot of the magnetically modulated photonic qubit output further quantifies that the R10 mutant exhibits a significantly higher lock-in amplitude than other rounds, demonstrating the most outstanding magnetic modulation performance (Figure 5b). Based on the dual analysis of time-domain waveforms and quantized amplitudes, we definitively selected the tenth-round mutant MagLOV 2 due to its optimal magnetic modulation performance. This variant, with its outstanding magnetic modulation fluorescence characteristics, provides a critical reporter unit choice for constructing highly sensitive biosensor systems for heavy metal ion detection. This selection robustly ensures the precise and efficient detection of heavy metal ions by this quantum biosensing system.

**Figure 5.**
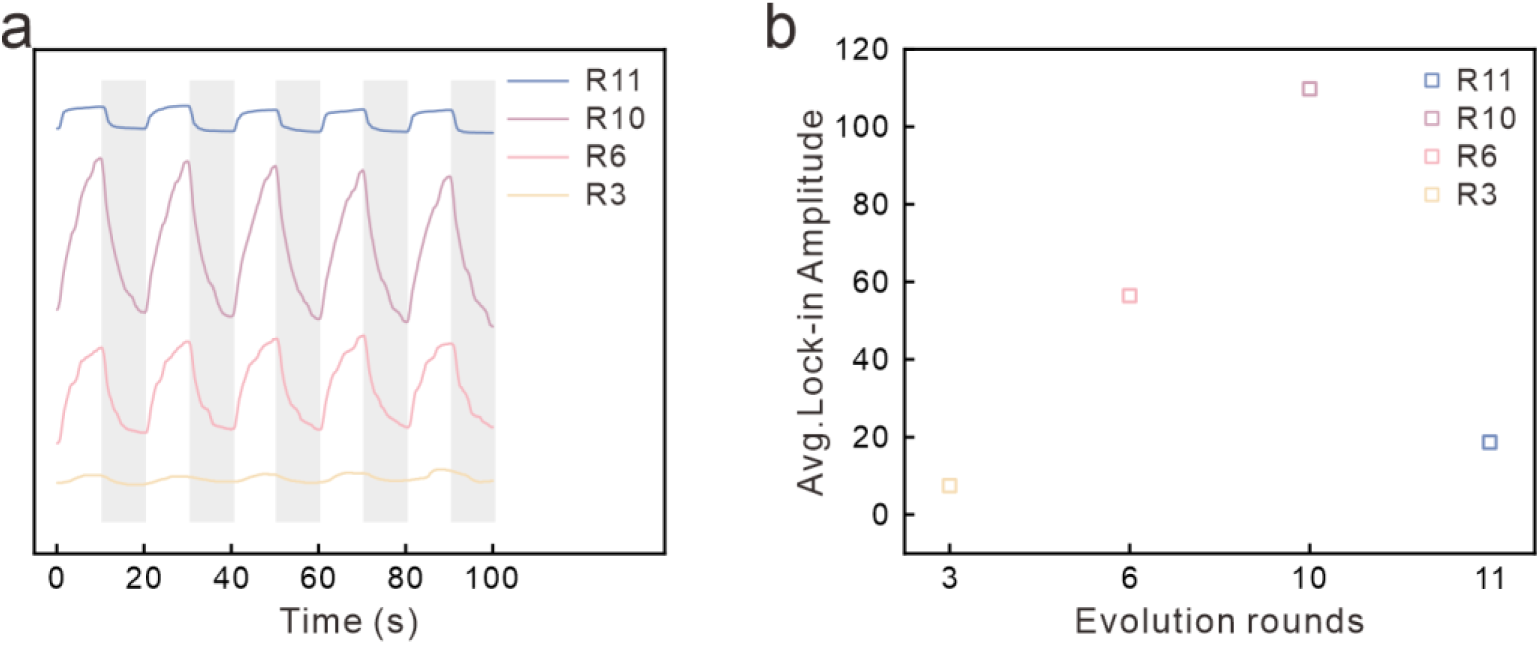
**a)** At a magnetic field strength of 10 mT, the time-domain profile of the magnetically modulated optical qubit of four representative MagLOV mutants from the directed evolution in BL21(DE3), and **b)** the lock-in amplitude output of the magnetically modulated optical qubit obtained via fast Fourier transform.

## 5. Conditional Optimization and Performance Appraisal of the pT7cadO−H MagLOV 2 Heavy Metal Assay System

pT7cadO−H MagLOV 2 heavy metal sensing system is constructed based on the CadC−T7 heavy metal response molecular mechanism (Figure 6a).^[37,38]^ Vector construction details are provided in S1.3. Its core response logic comprises three key steps: First, when heavy metal ions such as cadmium or lead are present in the environmental medium, these ions specifically stimulate the cadmium-responsive protein CadC. Thus, the binding between stimulated CadC and *cadO* triggers transcriptional activation of the *P*_T7_; Subsequently, the activated promoter drives the transcription and translation of the downstream reporter gene *MagLOV 2*, ultimately expressing the magnetically modulated fluorescent protein MagLOV 2; Finally, lock-in detection technology captures the periodic fluctuations of the MagLOV 2 photonic qubit signal. By calculating the SBR, quantum-level quantitative analysis of heavy metal ion concentrations is achieved.

**Figure 6.**
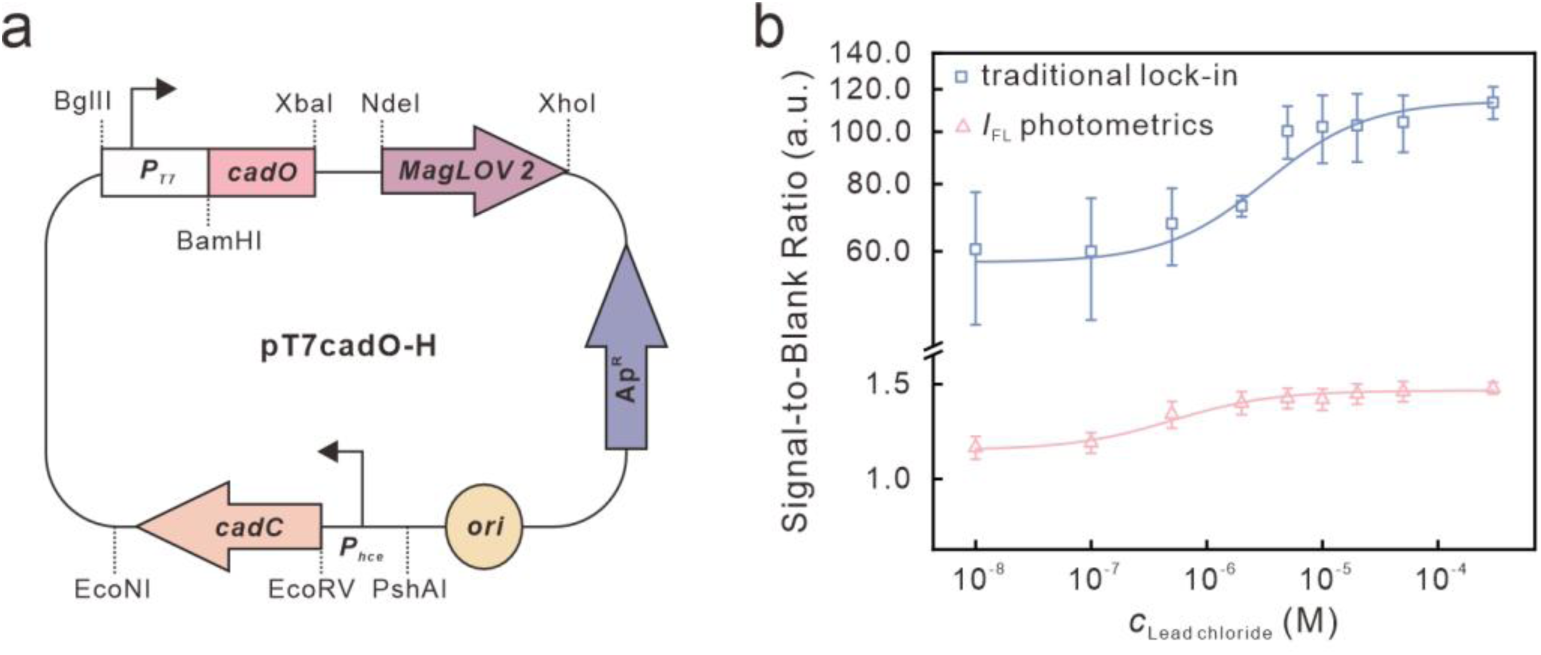
**a)** Schematic diagram of the pT7cadO−H MagLOV 2 recombinant plasmid design. **b)** Comparison of the lock-in amplitude and steady-state fluorescence SBR of pT7cadO1945−H MagLOV 2 detecting lead chloride at a magnetic field strength of 10 mT, fitted to a Langmuir isotherm calibration curve. The background represents the lock-in amplitude without lead chloride stimulation.

We conducted the initial experiment using lead chloride as the stimulus for 4 h. Heavy metal stimulation is shown in S1.6, while lock-in analysis and curve fitting are presented in S1.7. As shown in the experimental results (Figure 6b), under a constant magnetic field strength of 10 mT, the SBR of the magnetically modulated optical quantum bit lock-in analysis exhibited a steep dose-dependent response trend with increasing lead chloride concentration. Further fitting of the data to a Langmuir isotherm revealed a fitted curve with a coefficient of determination close to 1 (*R*^2^ = 0.942), indicating excellent positive correlation. In contrast, the SBR change observed with the traditional steady-state fluorescence method was gradual, demonstrating significantly lower sensitivity than the magnetically modulated optical quantum bit lock-in detection. This series of results fully demonstrates that the pT7cadO−H MagLOV 2 sensing system, through the complete response chain of heavy metal stimulation → stimulated CadC binding to *cadO* → MagLOV 2 expression → magnetic modulation of the optical quantum bit signal output, can achieve highly sensitive and highly specific detection of heavy metal ions. High-specificity detection of heavy metal ions. This lays a solid technical foundation for its practical applications in real-time monitoring of environmental heavy metal pollution and rapid screening of heavy metal residues in aquatic environments.

Magnetic modulation frequency and heavy metal stimulation duration serve as core parameters directly influencing the output efficiency of optical qubit signals. Their optimization is crucial for ensuring system performance. To further enhance the sensitivity, specificity, and statistical reliability of the pT7cadO−H MagLOV 2 system for detecting heavy metals (represented by lead chloride), we conducted targeted optimization experiments for these two parameters. All tests were performed at a magnetic field strength of 10 mT.

For magnetic modulation frequency optimization, we selected five frequency gradients: 0.030 Hz, 0.040 Hz, 0.050 Hz, 0.075 Hz, and 0.100 Hz. We measured the lock-in amplitude output of the magnetically modulated optical qubit at each frequency (Figure 7a). Quantitative results reveal significant variations in lock-in amplitude across frequencies: the amplitude peaks at 0.05 Hz within the experimental range. Deviations from 0.05 Hz - whether decreasing to 0.030 Hz or 0.040 Hz, or increasing to 0.075 Hz or 0.100 Hz - result in diminishing lock-in amplitudes to varying degrees. This result clearly indicates that 0.05 Hz is the optimal frequency parameter for magnetic modulation detection in this system. Selecting this frequency maximizes both the intensity of the magnetically modulated optical quantum bit signal and the detection sensitivity, providing precise parameter guidance for subsequent signal acquisition in heavy metal detection.

**Figure 7.**
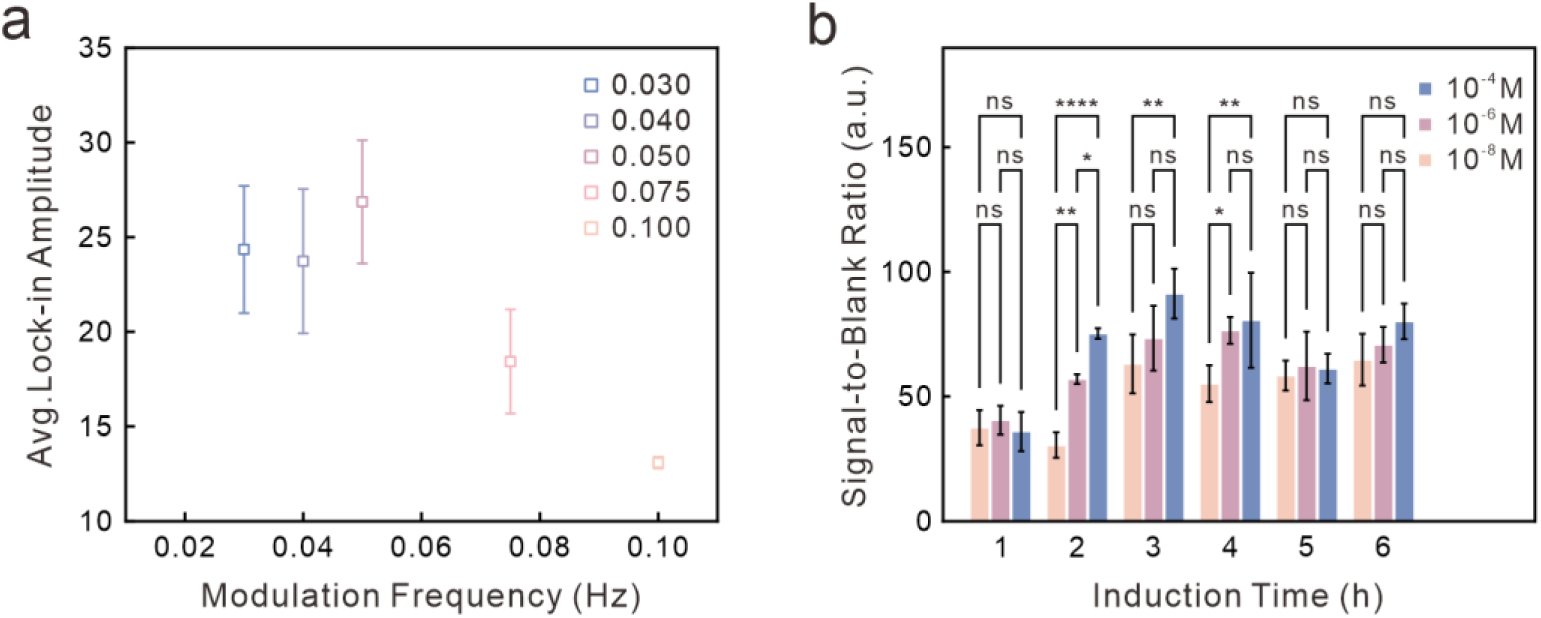
Optimized parameters of the pT7cadO1945−H MagLOV 2 system for detecting lead chloride. At a magnetic field strength of 10 mT: **a)** Lock-in amplitude plots of magnetically modulated optical qubits obtained via fast Fourier transform under different magnetic field frequencies; **b)** Intra-group significant differences in SBR of lock-in amplitude between lead chloride groups (orange, 10^−8^ M; purple, 10⁻⁶ M; blue, 10⁻⁴ M) and the no-lead chloride background group at different stimulation durations (1∼6 h, row factor). Statistical analysis employed two-way ANOVA (α = 0.05) combined with Tukey’s multiple comparison test, with "stimulation duration" as the row factor and "lead chloride concentration" as the column factor. This assessed SBR differences among concentration groups and the background group at the same stimulation duration. Significance notation: ns indicates *P* ≥ 0.05 (no significant difference), * indicates *P* < 0.05, ** indicates *P* < 0.01, *** indicates *P* < 0.001, **** indicates *P* < 0.0001. Data represent the mean of 3 parallel experiments, with error bars denoting standard deviation.

To optimize heavy metal exposure duration, we established six-time gradients (row factor): 1 h, 2 h, 3 h, 4 h, 5 h, and 6 h. Concentration gradients (column factor) comprised 10^−8^ M, 10^−6^ M, and 10^−4^ M lead chloride. Intra-group differences were analyzed using two-way ANOVA combined with Tukey’s multiple comparison test (Figure 7b). The two-way ANOVA revealed that both stimulation duration (F(5,36) = 21.66, *P* < 0.0001) and lead chloride concentration (*F*(2,36) = 20.09, *P* < 0.0001), and their interaction (*F*(10,36) = 2.905, *P* = 0.0091) significantly influenced the SBR. Stimulation duration contributed most to the total variance (50.71%), emerging as the primary factor regulating the detection signal. Tukey’s multiple comparison further confirmed that the 2-h stimulation group exhibited the most significant differences: 10^−8^ M *vs*. 10^−6^ M (*P* = 0.0032, **), 10^−8^ M *vs*. 10^−4^ M (*P* < 0.0001, ****), 10^−6^ M *vs*. 10^−4^ M (*P* = 0.0489, *). These concentrations exhibited optimal SBR discrimination. In contrast, the 1-h group showed no significant differences between concentrations due to insufficient signal accumulation (*P* > 0.05, ns). Within the 3-h group, only 10^−8^ M *vs*. 10^−4^ M was significant (*P* = 0.0016, **), with no statistical differences in other comparisons. In the 4-h group, 10^−8^ M *vs*. 10^−6^ M (*P* = 0.0188, *) and 10^−8^ M *vs*. 10^−4^ M (*P* = 0.0046, **) showed significant differences, but 10^−6^ M *vs*. 10^−4^ M showed no difference; No significant differences were observed in any comparisons within the 5-h and 6-h groups (*P* > 0.05, ns), likely due to cellular metabolic saturation and plateauing protein expression. These statistical results conclusively confirm that 2 hours represents the optimal lead chloride stimulation duration, ensuring sufficient signal accumulation while enabling efficient differentiation between concentrations and guaranteeing the statistical validity of detection results.

After the Condition Optimization, to validate the applicability of the pT7cadO−H MagLOV 2 system for detecting multiple heavy metal compounds and to clarify the performance differences between various detection methods, we conducted concentration gradient detection experiments on three target compounds: lead chloride, lead nitrate, and cadmium chloride, using three detection methods, i.e., MagLOV 2-based lock-in analysis, MagLOV 2-based *I*_FL_ (*I*_FL_MagLOV 2_) photometrics, and EGFP-based *I*_FL_ (*I*_FL_EGFP_) photometrics. Figure 8 presents the SBR Langmuir isotherm calibration curves for the three heavy metals across the 10^−11^ to 10^−4^ M concentration range, under a magnetic field strength of 10 mT. The blue curve representing lock-in analysis exhibits a pronounced dose-dependent response in the concentration gradients of lead chloride (Figure 8a), lead nitrate (Figure 8b), and cadmium chloride (Figure 8c), maintaining clear signal discrimination in the 10^−8^ to 10^−4^ M range. In contrast, the pink curve representing *I*_FL_MagLOV 2_ and the orange curve for *I*_FL_EGFP_ exhibited relatively flat SBRs with indistinct signal differences at low concentrations, intuitively demonstrating the sensitivity advantage of lock-in analysis in low-concentration heavy metal detection. Regarding detection limits, lock-in analysis achieved nanomolar detection limits for all three heavy metal compounds (lead chloride: 9.9 nM, lead nitrate: 4.8 nM, cadmium chloride: 2.4 nM). This represents a 1∼2 order-of-magnitude improvement in sensitivity compared to *I*_FL_EGFP_ (1.35∼1.48 μM) and the micromolar detection limits of *I*_FL_MagLOV 2_ (2.76∼4.33 μM). Regarding detection accuracy, lock-in analysis consistently achieved *R*^2^ values exceeding 0.95. Specifically, *R*^2^ values for lead nitrate (0.995) and cadmium chloride (0.998) approached 1, indicating excellent fitting performance. In contrast, the *R*^2^ values for *I*_FL_MagLOV 2_ were relatively lower (0.874∼0.983), while those for *I*_FL_EGFP_ exhibited fluctuations (0.902∼0.970). The pT7cadO−H MagLOV 2 system demonstrates detection capabilities for multiple heavy metal compounds including lead chloride and cadmium chloride. Compared to *I*_FL_ photometrics, MagLOV 2-based lock-in analysis demonstrates significant advantages in detection sensitivity and quantitative accuracy, providing critical technical support for high-sensitivity quantum monitoring of diverse heavy metal pollutants.

**Figure 8.**
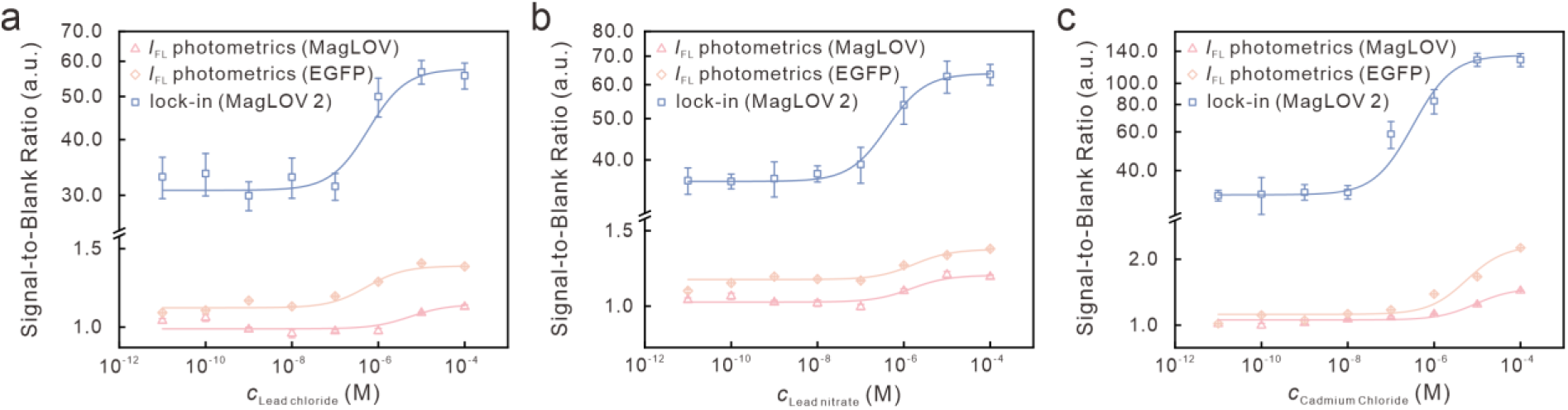
pT7cadO−H MagLOV 2 (cadO1945, cadO640) multianalyte detection capability and comparison of multimodal measurement methods. At a magnetic field strength of 10 mT, Langmuir isotherm calibration curves fitted for SBRs at different target stimulus concentrations: **a)** lead chloride (10^−11^∼10^−4^ M); **b)** lead nitrate (10^−11^∼10^−4^ M); and **c)** Cadmium chloride (10^−11^∼10^−4^ M). Blue: MagLOV 2-based lock-in analysis; Pink: MagLOV 2-based *I*_FL_ photometrics; Orange: EGFP-based *I*_FL_ photometrics.

## 6. DISCUSSION

This study successfully constructed a magnetically modulated optical quantum bit biosensing system based on MagLOV 2 and CadC−T7 gene circuit, demonstrating significant quantum performance advantages in the highly sensitive detection of multiple heavy metal ions. Its value can be analyzed from multiple dimensions: Compared to traditional fluorescent protein reporter units, MagLOV 2’s magnetically modulated quantum properties overcome the background interference limitations inherent in fluorescence detection. This represents the core rationale for its significantly enhanced sensitivity in low-concentration heavy metal detection. System optimization of magnetic modulation frequency, heavy metal stimulation duration, and *E. coli* detection concentration establishes the operational foundation for quantum performance. A modulation frequency of 0.05 Hz achieves the optimal balance between quantum signal intensity and detection efficiency. A 2-h heavy metal stimulation period ensures sufficient MagLOV 2 protein expression for effective signal accumulation while preventing cellular metabolic saturation or signal decay. Maintaining *E. coli* concentration at OD_600_ = 1 effectively avoids optical scattering/absorption quenching of fluorescence caused by high-turbidity bacterial cultures, guaranteeing quantum bit signal stability. This bio-quantum system achieves nanomolar detection limits for lead chloride, lead nitrate, and cadmium chloride, with Langmuir fitting *R*^2^ > 0.95. Compared to traditional fluorometric methods based on EGFP and MagLOV2, magnetic modulation quantum bit lock-in analysis enhances sensitivity by 1∼2 orders of magnitude. This advantage renders it irreplaceable in environmental trace heavy metal detection scenarios. For instance, trace cadmium and lead ions in actual water bodies often exist at nanomolar levels, which traditional methods may miss. This quantum biosensing system enables precise quantification, demonstrating dual advantages of high sensitivity and high quantitative accuracy, effectively overcoming the limitations of high background interference and limited sensitivity in conventional fluorescent sensors.

Although this system demonstrates outstanding performance in heavy metal detection, key limitations persist in practical application expansion that require targeted breakthroughs: First, detection coverage remains limited to a single target, currently developed only for a few heavy metal ions. Its reliance on the heavy metal - cadmium specific binding mechanism fails to address screening demands for heavy metal-organic pollutants mixed contamination in real-world environments; Second, its resistance to interference from complex sample matrices is weak. Existing performance validation is primarily based on ideal buffers or pure bacterial solutions. Components in real-world environmental samples (e.g., surface water, soil extracts), such as humic substances and high-salt content, may interfere with detection accuracy through optical scattering that disrupts magnetically modulated signals or by competing for binding with *cadO* elements. Furthermore, no suitable pretreatment protocols for complex samples have been established. Third, signal interpretation remains unidimensional, relying solely on the SBR of magnetically modulated optical qubits for pollutant concentration quantification.

This approach fails to correlate with toxicological effects (e.g., toxicity differences among heavy metals, cumulative effects of long-term exposure), precluding integrated concentration-ecological risk assessments and thus falling short of meeting deeper environmental monitoring requirements.

To advance the biosensor’s translational utility and address its current limitations, three key directions for optimization are proposed:

1. The scope of detectable analytes and the breadth of specificity could be substantially expanded through the integration of organic pollutant-responsive transcription regulatory modules including per- and polyfluoroalkyl substances (PFCs)-targeted protein pfc−DEF ^[39]^ and 2,4-dinitrotoluene (DNTs)-responsive promoters *azoR* and *yqjF*.^[40,41]^ Leveraging these elements, multianalyte responsive plasmid libraries could be constructed to enable high-resolution differentiation and concurrent detection of co-occurring pollutants in mixed contamination scenarios, a critical capability for real-world environmental monitoring.
2. Compatibility with complex environmental matrices should be enhanced via the development of solid-phase extraction (SPE)-based sample pretreatment workflows and matrix purification protocols, which would mitigate interference from matrix components. Concurrently, tuning of magnetic modulation parameters would bolster system robustness across a broad range of pH values and ionic strengths, with benchmarking of performance mandated in authentic environmental matrices (including real-world surface water samples and soil extracts) to validate field applicability.
3. The dimensionality of signal interpretation could be broadened by establishing a concentration-toxicity correlation assessment framework. This would involve integrating complementary readouts into the biosensing platform - specifically, bacterial metabolic activity (monitored using fluorescent metabolic probes) and DNA damage (tracked via DNA repair protein biomarkers) - thereby extending the system’s utility from mere quantitative analyte detection to comprehensive ecological risk evaluation.

Building on the aforementioned refinements, and capitalizing on the quantum magnetic modulation properties and *in situ* expression capabilities of MagLOV 2, five synergistic and transformative functional upgrades can be realized via the following innovative strategies, which position the platform at the interface of quantum sensing and synthetic biology:

**1)** A multichannel photonic qubit detection system integrating wavelength-resolved magnetically modulated fluorescent proteins should be developed. By engineering a recombinant plasmid library encoding MagLOV 2 (green-emissive, heavy metal-responsive), mScarlet (red-emissive, per- and polyfluoroalkyl substance (PFC)-responsive)^[42]^ and yellow fluorescent protein (YFP, yellow-emissive, 2,4-dinitrotoluene (DNT)-responsive)^[43]^ - each fused to analyte-specific regulatory elements - coupling with a multichannel optical qubit lock-in detector would enable the deconvolution of orthogonal wavelength signals. This setup facilitates simultaneous identification and quantification of multiple co-occurring pollutants, thereby drastically boosting detection throughput and resolution for complex environmental matrices.
**2)** Harnessing MagLOV’s microwave (radiofrequency, RF)-modulated spin relaxation mechanism,^[36]^ synergistic optimization of microwave-magnetic field parameter regimes could unlock parallel detection of diverse biological targets, including DNA lesions and pathogen-specific gene sequences. Exploiting the heightened sensitivity of microwave-modulated spin relaxation, *in situ* cellular microenvironment sensors could be engineered to quantify local paramagnetic species concentrations and pH gradients. When combined with magnetic gradient imaging to mitigate scattering interference from biological tissues, this technology enables high-resolution spatial mapping of MagLOV-labeled DNA dynamics *in vivo*, bridging quantum sensing and live-cell imaging.
**3)** Organelle-targeted quantum detection modalities can be established by fusing MagLOV 2 and homologous fluorescent qubits to subcellular localization tags, including mitochondrial targeting sequences (MTS)^[44]^ and endoplasmic reticulum retention signals (KDEL).^[45]^ The resultant subcellularly localized magnetically modulated photonic qubit reporters, paired with high-resolution confocal magnetically modulated imaging, support real-time tracking of organelle-resident probe dynamics at the quantum scale, furnishing unprecedented visualization tools for subcellular cell biology research.
**4)** *In vivo* biological stimulus-sensing models can be constructed by integrating the magnetically modulated optical qubit system into tractable model organisms, from *E. coli* to zebrafish embryos. Monitoring fluctuations in quantum optical signals would decode biological responses to key environmental cues (e.g., analyte concentration, pH shifts, thermal perturbations), providing a quantum-enabled framework for mechanistic investigations of ecological stress responses and advancing the quantitative underpinnings of ecological risk assessment.
**5)** Leveraging native DNA-sensing machineries in live bacterial chassis,^[46]^ *Acinetobacter baylyi* and *Bacillus subtilis* can be repurposed to host stably genomically integrated MagLOV 2 (via homologous recombination). Functionalization of this system, either by fusing the DNA glycosylase OGG1 to quantify oxidative DNA damage, or by engineering a MagLOV 2−dCas9 CRISPR complex for target DNA tracking, would enable multimodal biosensing. Coupling with multi-wavelength qubit readouts and anti-interference algorithm optimization, this platform could be translated to diverse high-stakes applications: clinical detection of DNA damage-associated diseases and pathogenic infections, food safety screening for foodborne pathogens, and environmental monitoring of pollutant-induced genotoxicity.

In summary, the MagLOV 2 magnetically modulated optical qubit biosensing platform delivers a quantum-scale solution for ultra-trace contaminant detection and live-system biological sensing. Its modular design unlocks transformative potential across environmental monitoring, cell biology and translational diagnostics, laying the groundwork for next-generation quantum-enabled biosensing in diverse ecological and biomedical scenarios.

## Supporting information

Suppporting Information

## SUPPORTING INFORMATION

Supporting Information is available from the bioRxiv or from the author.

## ACKNOWLEDGEMENTS

This work was financially supported by National Natural Science Foundation of China (Grant Nos. 22374076, 22204077, 22504061), Natural Science Foundation of Jiangsu Province (BK20231455), Fundamental Research Funds for the Central Universities (2025201001, 30924010201), Open Research Program of National Major Scientific and Technological Infrastructure for Translational Medicine (TMSK−2024−115), State Key Laboratory for Analytical Chemistry for Life Science (SKLACLS2402), and Postgraduate Research & Practice Innovation Program of Jiangsu Province (KYCX25_0763).

## DECLARATIONS

### Unique biological materials

All biological materials (bacterial cells, plasmids) are available upon request.

### Author contribution

- Wenhao Shan − cloning plasmids and strains strategy, protein expression, experimental measurements, analysis, writting.
- Fujin Lv − sample preparation, protein expression, purification and quantification.
- Wei Liu − magnetic field modulation device design, lock-in analysis strategies and algorithm development.
- Bin Li − microscope platform development, design and development of imaging experiments.
- Yingying Wang − algorithm development and optimization.
- Xianli Gong − magnetic field modulation device design and construction.
- Fulin Zhu − protein expression advice.
- Bingchen Zeng − sample preparation, experimental measurements.
- Tiantian Man − manuscript figure advice.
- Qiuyue Wu − manuscript writing advice.
- Shengyuan Deng − project co-ordination and conceptualisation, supervision.

