## Supplementary material for "Stimuli−Expressed Protein Qubits for Quantum Quantification of Heavy Metals in Natural Water": Suppporting Information

### TABLE OF CONTENTS


#### 1. Supplementary Experimental Details

Strains S−3

Enzymes and Chemicals S−3

Plasmid Construction and Sequence S−3

MagLOV 2 Expression and Purification S−4

Magnetic Field Generation and Signal Measurement S−5

Heavy Metal Exposure Treatment of MagLOV 2 Bacterial Culture S−6

#### 2. Supplementary Notes

Opto-Qubit Lock-in Quantitation and Curve Fitting S−7

#### 3. Supplementary Code

Lock-in Amplitude Analysis Code S−8

#### References

### 1. Supplementary Experimental Details

#### Strains

The strains used in this study are Escherichia coli DH5α and BL21(DE3). DH5α is employed for plasmid amplification, while BL21(DE3) is used for gene expression.

#### Enzymes and Chemicals

The kits used in the molecular cloning experiments of this study were purchased from Tiangen Biotech Co., Ltd. DNA polymerase and T4 ligase were acquired from Novazene Biotech Co., Ltd. Restriction enzymes for plasmid construction were purchased from New England Biolabs. All other reagents were sourced from Shanghai Bide Pharmaceutical Technology Co., Ltd., McLean Biotech Co., Ltd., Shanghai Sangon Biotech Co., Ltd., and Beijing Kangwei Century Biotechnology Co., Ltd.

#### Plasmid Construction and Sequence

The MagLOV 2 gene expression vector is the prokaryotic expression vector pET-28a(+), while the CadC−T7 gene circuit vector is the prokaryotic expression vector pET-21a(+). *MagLOV 2*, *cadO*, *cadC*, $P_{T7}$ and $P_{hce}$ were synthesized by Sangon Biotech (Shanghai) Co., Ltd.[1,2] The 6×His-*MagLOV 2* gene was inserted into *NcoI*/*XhoI*-digested pET−28a(+), yielding pET-28a(+)-MagLOV 2. The $P_{T7}$-*cadO*-*MagLOV 2* was inserted into *BglII*/*XhoI*-digested pET-21a(+) to replace the *lac* operon, yielding pT7cadO MagLOV 2. Subsequently, the $P_{hce}$ -*cadC* was inserted into pT7cadO MagLOV 2 pretreated with *PshAI*/*EcoNI* to replace *lacI*, yielding pT7cadO−H MagLOV 2; *BamHI*/*XbaI* sites were designed at both ends of *cadO*, *NdeI*/*XhoI* sites at both ends of *MagLOV 2*, and *EcoRV*/*EcoNI* sites at both ends of *cadC* to facilitate replacement of heavy metal-specific response elements and reporter genes. Successful plasmid construction was verified by DNA sequencing.

**MagLOV 2 coding sequence:**

ATGCTTGCTACTACTTTAGAACGTATAGAGAAAAACTTCGTGATCACGGACCCGAGACTACCTGAC
AACCCTATAATTTTTGCAAGTGACTCATTCCTTCAGTTGACTGAGTATTCTAGGGAAGAGATTCTA
GGGTGGAATCCTAGATTCTTGCAAGGACCAGAAACTGACCGTGCCACTGTGAGGAAAATCAGGGA
TGCGATCGACAACCAAACCGAGGTGACAGTGCAGCTAATAAATTACACTAAATCTGGCAAGAAGT
TCTGGAACCTACTTCATGTGCAACCCATGAGAGACCAAAAAGGAGACGTACAGTACTTCATAGGG
GTAAAGTTGGATGGTACTGAGCATGTTAGAGACGCGGCAGGTCGTGAACGTGTTATGTTAATAAA
AAAGACCGCTGAAAACATAATGGAAGCGGCAAAGGAGTTGTAA

**Coding sequence of heavy metal response elements:**

***cadO640*:** AATAATCAAATGATTGTTTGAGTATG

***cadO1945*:** TATATTCAAACATACACTTGAATAAA

***cadC640*:**

ATGAAACAGGATGACGTTTGCGAAGTTACCTGCGTTGATGAAGAAAAAGTTCGTCGTGTTAAAGA
AAGCGTTAAACAGCAGAACACCCTGGCGGTTAGCCAAATCTTCAAAGCGCTGAGCGATGATACCC
GTGTTAAAATCACCTTCAGCCTGTACGAAGAAGAAGGCCTGTGCGTTTGCGATGTTGCGAACATCG
TTGGCTGCACCACCGCGACCGCGAGCCACCACCTGCGTCTGCTGCGTAACATGGGCCTGGCGAAAT
ACCGTAAAGAAGGCAAACTGGTTTTCTACAGCCTGGATGATGACCACGTTCGTCAGCTGATCCAGA
TCGCGTTCGCGCACCAGAAAGAAGTTGAAAACTACGAATAA

***cadC1945*:**

ATGAGCAAAAAAGATACCTGTGACATTTATTGTTATGATGAAGCGAAAGTGAAACGCATTCAGGG
CGAAATGCAGAAAGAAGATATTAGCAGCGTGAGCCAGCTGTTTAAAGCGCTGGCGGACGAAAATC
GTGCGAAAATTAGCTACGCGCTGTGCCAGGATGATGAACTGTGTGTGTGTGATGTGGCCAACATTA
TTGGCGCGACCGTGGCGACCACCAGCCATCATCTGCGTACCCTGCATAAACAGGGCATTGTGAAAT
ACCGCAAAGAAGGCAAACTGGCGTTTTACAGCCTGGATGATGAGCATATTCGCCAGCTGATGGTG
ATTGCGCTGACCCACAAAAAAGAAATGAAAGTTAACGTGTAA

#### MagLOV 2 Expression and Purification

The recombinant plasmid pET-28a(+)-MagLOV 2 was transformed into BL21(DE3) cells and sequenced for verification. A seed culture was grown in 10 mL LB medium containing 100 μM kanamycin. After overnight incubation at 37 °C, the seed culture was then diluted 1:100 into 100 mL LB medium containing 100 μM kanamycin and cultured continuously at 37 °C with shaking. At an OD600 of 0.6~0.8, induction was performed with 1 mM IPTG, followed by incubation in the dark at 27 °C for 16 h. The overnight−induced cells were collected by centrifugation at 4500× g for 10 min. The pellet was resuspended in Buffer A and sonicated. After centrifugation at 16,000× g for 45 min at 4°C, the cell-free lysate was loaded onto a Ni-NTA column. The column was washed with Buffer A containing 35 mM imidazole and eluted with Buffer A containing 250 mM imidazole. Buffer A dialysis was performed to remove imidazole. Purified protein was concentrated and stored at −80 °C. Protein purification removed most contaminating proteins, with MagLOV (molecular weight 16.7 kDa) as the major protein product (Figure S1). Protein concentration was determined using Nano-300 (ALLSHENG, China).

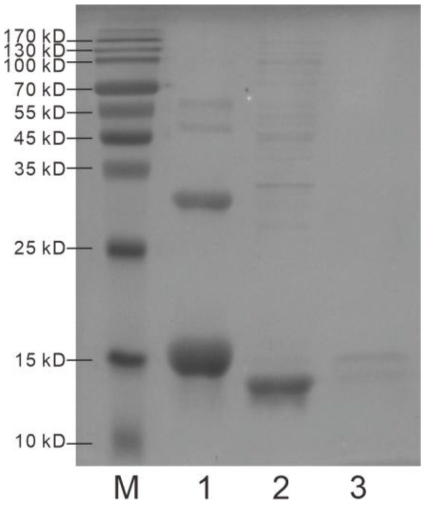


**Figure S1.** SDS-PAGE analysis of MagLOV 2 expression and purification. Lane M: protein marker; Lane 1: concentrated protein sample; Lane 2: cell lysate supernatant; Lane 3: protein purification eluate.

#### Magnetic Field Generation and Signal Measurement

This periodic magnetic field modulation device adopts a core architecture of "signal generation-amplification drive-power supply assurance-magnetic field output". It utilizes the signal generator FY6900 (0~100 MHz, 0~24 Vpp, FeelElec, China), PWM signal amplifier YF-44 (0~10 KHz, 3~5 V, YUEYU, China), power adapter KJS-1510 (3~24 V, KJS, China), and the core-type electromagnet KK-P50/30 (Kaka Electric, China). This system achieves precisely controllable periodic magnetic field output (Figure S2). The signal generator serves as the "signal source" for magnetic field modulation, outputting raw PWM signals with adjustable frequency and duty cycle. It directly determines core parameters such as the modulation period and waveform characteristics of the periodic magnetic field. The PWM signal amplifier receives the output signal from the signal generator and amplifies the weak PWM signal, enhancing its drive capability to ensure stable response to modulation commands from subsequent execution components. The power adapter provides stable, matched operating power to the entire system, supplying the signal generator, PWM signal amplifier, and iron-core electromagnet to guarantee the stability and consistency of each component's operation. The iron-core electromagnet serves as the core execution component for magnetic field output. It receives amplified PWM drive signals and generates corresponding periodic magnetic fields through periodic current changes. Its iron-core structure effectively enhances magnetic field strength and focusing effects. The electromagnet's magnetic field strength is measured by the GM-55 Gmeter (0~2000 mT, Tindun Industry, China). The overall workflow involves the signal generator outputting a PWM signal with preset parameters. This signal is amplified by the PWM signal amplifier to drive the iron-core electromagnet, while the power adapter provides stable power supply throughout. Ultimately, the iron-core electromagnet outputs a periodic modulated magnetic field meeting the requirements, featuring a simple architecture and precise control. The acquisition and presentation of optical qubit signals rely on a spectrophotofluorometer (Light source: 150 W, excitation/emission: 200~900 nm, Lengguang Tech, China) captures and converts optical qubit signals in time–scan mode. After data transmission, key information including signal characteristics and temporal variations is displayed in real time on the host computer interface, enabling visual observation and data logging of optical qubit signals.

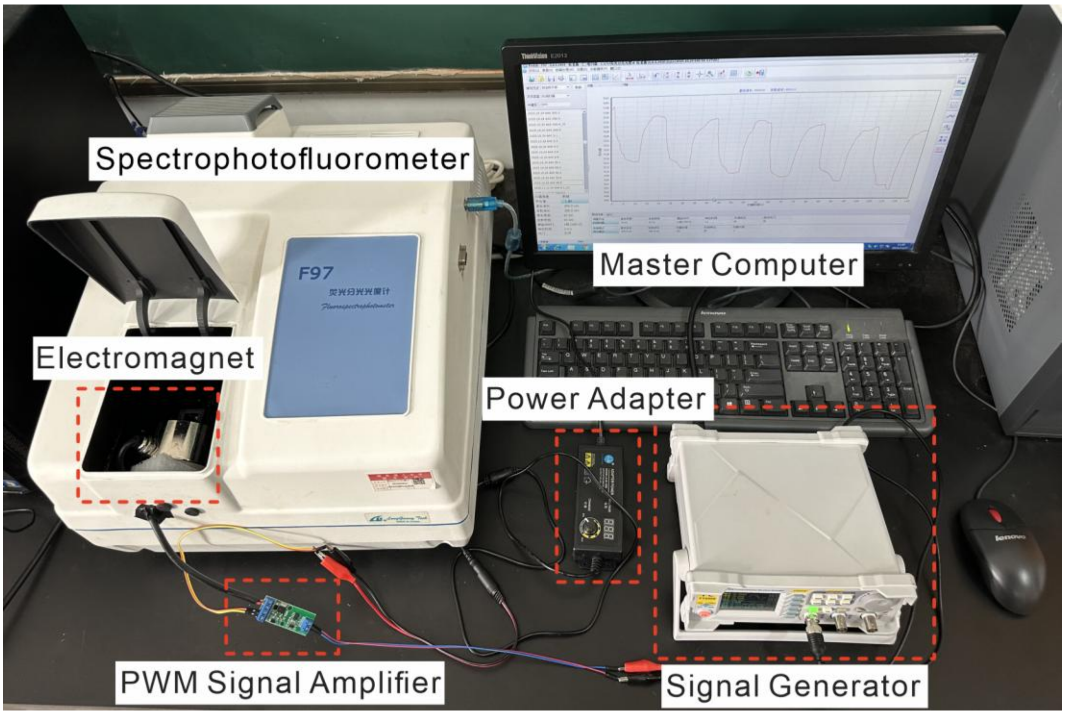


**Figure S2.** Magnetic field modulation and signal output detection prototype.

#### Heavy Metal Exposure Treatment of MagLOV 2 Bacterial Culture

The recombinant plasmid pT7cadO−H MagLOV 2 was transformed into BL21(DE3) cells and sequenced for identification. A seed culture was grown in 10 mL LB medium containing 100 μM ampicillin. After overnight incubation at 37 °C, the seed culture was then expanded at a 1:100 ratio into 100 mL LB medium containing 100 μM ampicillin and cultured continuously at 37 °C with shaking. When the $OD_{600}$ reached 0.4~0.5, heavy metal compounds at various designed concentrations were added as stimuli. The cells were stimulated for 2 h at 37 °C in the dark. Overnight-induced cells were harvested by centrifugation at 4500× g for 10 min. The pellet was resuspended in PBS buffer to achieve an $OD_{600} \approx 1.0$, yielding the detection sample. Heavy metal compounds (cadmium chloride, lead chloride, lead nitrate) were dissolved in sterile water and sterilized by filtration.

#### 2. Supplementary Notes

##### Opto-Qubit Lock-in Quantitation and Curve Fitting

The amplitude of the FL signal fluctuation is determined through a lock-in algorithm. For each fluorescence detection of the target sample, the average fluorescence intensity is calculated based on the sampling data from the spectrophotofluorometer, i.e., spanning the time axis from the initial time $t_0 = 0$ to the termination time $t_N = N/F_S$ (where $N$ is the number of sampling points within a single cycle, and $F_S$ is the sampling frequency). After converting the modulation frequency ($F_M$) to angular frequency ($2\pi \cdot F_M$), the fluorescence intensity $I_{FL}$ at time $t_N$ is first multiplied by both the in-phase reference signal $\sin(2\pi \cdot F_M \cdot t_N)$ and the 90° out-of-phase signal $\cos(2\pi \cdot F_M \cdot t_N)$ to obtain $I_x$ and $I_y$, respectively, as follows:

$$I_x = I_{\mathrm{FL}} \cdot \sin(2\pi \cdot F_{\mathrm{M}} \cdot t_N) \quad \textbf{(S1)}$$

$$I_y = I_{\mathrm{FL}} \cdot \cos(2\pi \cdot F_{\mathrm{M}} \cdot t_N) \quad \textbf{(S2)}$$

By calculating the mean-value decomposition of the DC components of $I_x$ and $I_y$, denoted as $\overline{I_x}$ and $\overline{I_y}$ respectively, the lock-in amplitude ($A_{li}$) corresponding to $F_M$ can be expressed as:[3]

$$A_{\mathrm{li}} = \sqrt{\overline{I_x}^2 + \overline{I_y}^2} \quad \textbf{(S3)}$$

For faint yet genuine signals, even when obscured by spatially inhomogeneous or relatively stable background noise, the signal-to-noise ratio (SBR) remains a valid metric for assessing imaging quality, as it correlates with the "contrast" parameter of the target fluorophore. For modulated ($SBR_M$, derived from lock-in analysis) and demodulated ($SBR_{dM}$, derived from photometric measurements) scenarios, the formulae are as follows:

$$\mathrm{SBR_M} = \frac{mP_{\mathrm{S}}\sqrt{n}}{\sqrt{P_{\mathrm{B}}}} \quad \textbf{(S4)}$$

$$\mathrm{SBR_{dM}} = \frac{P_{\mathrm{S}}}{P_{\mathrm{B}}} \quad \textbf{(S5)}$$

In the equation, $P_S$ denotes the number of photons emitted during steady-state fluorescence, $P_B$ represents the background photon count, $m$ is the signal amplification factor, and $n$ indicates the number of photon counting chambers.[4] With appropriate values for $m$ and $n$, $SBR_M$ can achieve effective enhancement regardless of background noise intensity. Combining the definitions of fluorescence intensity $I_{FL}$ in Eq. S2 and S3, the SBR formulas of Eq. S4 and S5 are restructured as follows:

$$\mathrm{SBR_M} = \frac{mA_{\mathrm{li \cdot S}}\sqrt{N}}{\sqrt{A_{\mathrm{li \cdot B}}}} = 10 \cdot \frac{A_{\mathrm{li \cdot S}}}{\sqrt{A_{\mathrm{li \cdot B}}}} \quad \textbf{(S6)}$$

$$\mathrm{SBR_{dM}} = \frac{I_{\mathrm{FL \cdot S}}}{I_{\mathrm{FL \cdot B}}} \quad \textbf{(S7)}$$

$I_{FL \cdot S}$ and $I_{FL \cdot B}$ are set as the average fluorescence intensities for the heavy metal gradient addition group (target group) and the nonheavy metal addition group (background group), respectively; $A_{li \cdot S}$ and $A_{li \cdot B}$ are attributed to the locked amplitude and background of MagLOV 2, respectively. By default, $m$ and $N$ are set to 1 and 100, respectively. For simplicity, the combinations ($\mathrm{SBR_M}$, $\mathrm{SBR_{dM}}$) and ($A_{\mathrm{li \cdot S}}$, $A_{\mathrm{li \cdot B}}$) are generally referred to as SBR and $A_{\mathrm{li}}$ in the main body.

It can be inferred that when heavy metal target content is high, $I_{\mathrm{S}}$ reaches a plateau (i.e., MagLOV 2 expression saturates within the cell), allowing the relationship between SBR and heavy metal compound concentration ($c_{\text{heavy metal}}$) in Eq. S6 to be fitted as a Langmuir isotherm:[5]

$$\mathrm{SBR} = \frac{\mathrm{SBR_0} \cdot c_{\text{heavy metal}}}{K_{\mathrm{D}} + c_{\text{heavy metal}}} \tag{S8}$$

in which $K_{\mathrm{D}}$ represents the equilibrium dissociation constant. To determine the limit of detection (LOD), linear regression fits were performed between MagLOV 2's $A_{\mathrm{li \cdot S}}$ and $c_{\text{heavy metal}}$ under low-concentration heavy metal compound stimulation. Consequently, the LOD can be derived by dividing the background deviation ($\delta$, with at least three replicates) by the slope of the calibration curve within the low-concentration range:

$$\mathrm{LOD} = \frac{3\delta}{\mathrm{slope}} \tag{S9}$$

#### 3. Supplementary Code

##### Lock-in Amplitude Analysis Code

A custom Python script was used for quantitative analysis of spectrofluorometric data to extract lock-in amplitude and calculate SBR for characterizing magnetic protein fluorescence responses. The input CSV file contained parallel experimental data (Column C as DC reference; Columns I~N as six experimental groups), with 1000 fluorescence intensity points per column (10 Hz sampling rate). The workflow includes: **[1]** importing and validating data, then removing DC offset to isolate AC signals; **[2]** performing Fast Fourier Transform (FFT) and extracting the maximum normalized amplitude in the 0~16 Hz band as lock-in amplitude; **[3]** computing SBR via Eq. S6 (lock-in amplitude divided by the square root of (DC component/200), with DC value = 6.039041 for 1200 s conditions); **[4]** conducting statistical analysis (mean, SD, SEM) of SBR results; and **[5]** visualizing the reference group's raw data and 0–16 Hz spectrum for reliability verification. The code is shown below:

```
import pandas as pd

import numpy as np

import matplotlib.pyplot as plt

from scipy.fft import fft

import os    # Add os module for cross-platform path handling

# Configure matplotlib to display Chinese characters (retained for potential visualization; safe to remove if unused)

plt.rcParams['font.sans-serif'] = ['SimHei']    # Use SimHei font for Chinese characters

plt.rcParams['axes.unicode_minus'] = False        # Fix abnormal display of negative signs

# Define processing function: Extract the maximum normalized amplitude in the 0-16 Hz frequency range for a specified data column

def get_max_normalized_amp(data, sampling_rate):

    n = len(data)

    # Remove DC component to focus on AC signal components

    data_dc_removed = data - np.mean(data)

    # Perform Fast Fourier Transform (FFT) on DC-removed data

    fft_result = fft(data_dc_removed)

    # Calculate normalized amplitude spectrum (only retain positive frequency components)
```

```
    amplitude = np.abs(fft_result) / n * 2    # Double amplitude for positive frequencies
    amplitude = amplitude[:n//2]    # Keep only positive frequency half of the spectrum
    # Generate frequency axis corresponding to the amplitude spectrum
    freq_axis = np.linspace(0, sampling_rate/2, n//2, endpoint=False)
    # Filter amplitudes within the 0-16 Hz frequency range
    mask = (freq_axis >= 0) & (freq_axis <= 16)
    filtered_amp = amplitude[mask]
    # Return the maximum normalized amplitude in the 0-16 Hz range
    return np.max(filtered_amp)
# ----------------------
# Universal file path configuration (cross-platform & flexible)
# ----------------------
# Option 1: Relative path (RECOMMENDED for SI)
# Place your CSV file in the same folder as this script, then modify the filename below
file_name = "100.csv"    # Modify this to your actual CSV filename (e.g., "experiment_data.csv")
file_path = os.path.join(os.getcwd(), file_name)    # Auto-get current script folder + filename
# # Option 2: Manual input (flexible for users) - Uncomment if you need interactive path input
# file_path = input("Please enter the full path of your CSV file (e.g., C:/data/100.csv or
/Users/user/data/100.csv): ")
# Verify file exists (avoid runtime error)
if not os.path.exists(file_path):
    raise FileNotFoundError(f"File not found: {file_path}\nPlease check the filename/path or place the CSV
file in the script folder.")
# Read experimental data from CSV file
df = pd.read_csv(file_path)
# Basic experimental parameters
sampling_rate = 10    # Sampling rate: 10 Hz (samples per second)
data_length = 1000    # Number of data points (rows 1 to 1001, indexed as 0 to 1000)
```

```
# ----------------------
# Process the first experiment (Column C, index 2) - DC component reference
# ----------------------
c_data = df.iloc[0:data_length, 2].values    # Extract data from Column C
# dc_component = get_max_normalized_amp(c_data, sampling_rate)    # Calculate DC component reference
# print(f"First experiment (Column C) - Max normalized amplitude (DC component): {dc_component:.6f}")
# ----------------------
# Process subsequent 6 experiments (Columns I-N, indices 8 to 13)
# ----------------------
experiment_results = []
columns = [8, 9, 10, 11, 12, 13]    # Indices of target columns (I to N)
col_names = ['I', 'J', 'K', 'L', 'M','N']    # Corresponding column labels
for i, col_idx in enumerate(columns):
    # Extract data from the current column (rows 0 to 999)
    data = df.iloc[0:data_length, col_idx].values
    # Calculate maximum normalized amplitude (0-16 Hz) for current experiment
    max_amp = get_max_normalized_amp(data, sampling_rate)
    # Compute experimental result using the formula: normalized amplitude / sqrt(DC component / 200)
    # Note: DC component value (6.039041) is substituted for 1200s experimental condition
    result = max_amp / np.sqrt(6.039041 / 200)
    experiment_results.append(result)
    # Print detailed results for each experiment
    print(f"Experiment {i+2} (Column {col_names[i]}): Max normalized amplitude = {max_amp:.6f}, 
Calculated result = {result:.6f}")
# ----------------------
# Calculate statistical metrics for experimental results
# ----------------------
mean_value = np.mean(experiment_results)                    # Arithmetic mean of results
```

```
std_value = np.std(experiment_results)                    # Standard deviation (measures data dispersion)

sem_value = std_value / np.sqrt(len(experiment_results))    # Standard error of the mean (measures mean
reliability)

# Print statistical summary

print("\n===== Statistical Results =====")

print(f"Mean value: {mean_value:.6f}")

print(f"Standard deviation (data dispersion): {std_value:.6f}")

print(f"Standard error of the mean (mean reliability): {sem_value:.6f}")

# ----------------------

# Visualization module (fully uncommented for direct execution)

# ----------------------

# Extract data for plotting (Column C as reference)

x_data = df.iloc[0:data_length, 1].values    # X values from Column B

y_data = c_data    # Y values from Column C

# Plot raw data of the first experiment

plt.figure(figsize=(12, 8))

plt.subplot(2, 1, 1)

plt.plot(x_data, y_data, linewidth=0.8)

plt.title('First Experiment (Column C) - Raw Data')

plt.xlabel('X Value')

plt.ylabel('Y Value')

plt.grid(alpha=0.3)

# Calculate spectrum for visualization

n = len(y_data)

y_dc_removed = y_data - np.mean(y_data)

y_fft = fft(y_dc_removed)

amplitude = np.abs(y_fft) / n

amplitude = amplitude[:n//2]
```

```
freq_axis = np.linspace(0, sampling_rate/2, n//2, endpoint=False)
mask = (freq_axis >= 0) & (freq_axis <= 16)
filtered_freq = freq_axis[mask]
filtered_amp = amplitude[mask]
max_amp_idx = np.argmax(filtered_amp)
max_freq = filtered_freq[max_amp_idx]
# Plot frequency spectrum (0-16 Hz)
plt.subplot(2, 1, 2)
plt.plot(filtered_freq, filtered_amp, linewidth=1.0)
plt.scatter(max_freq, filtered_amp[max_amp_idx], color='red', label=f'Peak frequency: {max_freq:.4f} Hz')
plt.title('First Experiment (Column C) - Frequency Spectrum (0-16 Hz)')
plt.xlabel('Frequency (Hz)')
plt.ylabel('Normalized Amplitude')
plt.grid(alpha=0.3)
plt.legend()
plt.tight_layout()
plt.show()
```
